# Yield performance of chromosomally engineered durum wheat-*Thinopyrum ponticum* recombinant lines in a range of contrasting rain-fed environments across three countries

**DOI:** 10.1101/313825

**Authors:** Ljiljana Kuzmanović, Roberto Ruggeri, Jason A. Able, Filippo M. Bassi, Marco Maccaferri, Roberto Tuberosa, Pasquale De Vita, Francesco Rossini, Carla Ceoloni

**Affiliations:** Department of Agricultural and Forest Sciences (DAFNE), University of Tuscia, Viterbo, Italy; School of Agriculture, Food and Wine, Waite Research Institute, The University of Adelaide, Glen Osmond, Australia; ICARDA, Biodiversity and Integrated Gene Management, Rabat Institutes, Rabat, Morocco; Department of Agricultural Sciences, University of Bologna, Bologna, Italy; CREA-CI Cereal Research Centre for Cereal and Industrial Crops, Foggia, Italy

**Keywords:** Alien introgression, Chromosome engineering, Grain number, Tiller number, Wheat breeding

## Abstract

Introgressions of *Thinopyrum ponticum* 7AgL chromosome segments, spanning 23%, 28% and 40% of the distal end of durum wheat 7AL arm, were previously shown to contain multiple beneficial gene(s)/QTL for yield-related traits, in addition to effective disease resistance (*Lr19, Sr25*) and quality (*Yp*) genes. In the present study, durum wheat near isogenic recombinant lines (NIRLs), harbouring each of the three introgressions, were included for the first time in multi-location field trials, to evaluate general and environment-specific effects of the alien chromatin on 26 yield-related traits. The results from nine different trials across contrasting environments of Italy, Morocco and South Australia over four years revealed that the overall impact of 7AgL introgressions into the tetraploid wheat background did not imply, except in one environment, major yield penalty. The comprehensive effect of the three 7AgL segments on individual yield-contributing traits, resulted in significant increases of biomass m^−2^ (+9%), spike number m^−2^ (+13%), grain number m^−2^ (+11%) and spikelet^−1^ (+8%), but also in a general, significant decrease of grain weight (−8%). When the separate NIRLs were analysed, each of the three 7AgL segments turned out to be associated with variation of specific yield components. The effects of the 40%-long segment proved to be the most stably expressed across environments and involved significant increases of spike and grain number m^−2^ (13% and 15%, respectively), grain number spike^−1^ (10%) and spike fertility index (46%), though accompanied by a significant decrease in thousand grain weight (−23%). In spite of this trade-off between grain number and grain weight, their interplay was such that in four trials, including dryer environments, a grain yield advantage was observed. This evidence, and comparison with the two other NIRLs, substantiates the hypothesized existence of major gene(s)/QTL for grain number in the most proximal 28-40% 7AgL region, exclusive to the 40%-long 7AgL introgression. The present study represents an important validation of the use of chromosomally engineered genetic stocks for durum wheat improvement, targeting not only disease resistance and quality traits but also relevant yield components.

## Introduction

Durum wheat (*Triticum durum* var. *durum*, 2n = 4x = 28, genomes AB) is cultivated on approximately 8% of the world’s wheat area, and is economically important in the Mediterranean basin, North America’s Great Plains and Mexico, as well as Australia, Russia, Kazakhstan, India, Ethiopia and Argentina (Bassi and Sanchez-Garcia, 2017; Royo et al., 2014). Due to the socio-economic challenges emerging from the rapid human population increase across the world, breeding for continual improvement in yield potential (among other traits) is a priority for durum wheat breeders. Given that wheat provides calories for about 20% of human nutrition (FAO, 2013), current yield increases of not more than 1% year^−1^ for both durum and bread wheat, will be insufficient to meet the food demand in the imminent future (Fischer and Edmaedas, 2010; Ray et al., 2013). Further, with ongoing climate changes, especially temperature extremes and water shortages, already determining a progressive displacement of durum wheat cultivation from traditional areas (Ceoloni et al., 2014a; Habash et al., 2009), the goal of significantly increasing yield potential is even more challenging. An approximate 6% loss in global wheat yield was recently predicted for each degree-Celsius increase in temperature (Liu et al., 2016; Zhao et al., 2017). Yield reduction will depend on the interaction of temperature with other limiting factors, such as rainfall, CO_2_ emissions, and nitrogen supply, thus requiring environment-specific breeding strategies to select for highly adapted genotypes (Elía et al., 2018; Fleury et al., 2010; Luo et al., 2005; Tricker et al., 2018; Zhao et al., 2017). Consequently, for a particularly complex trait such as yield, understanding and dissecting the multi-layered genotype × environment interaction in multi-environment trials is a crucial step towards closing the gap between the actual and attainable yields, particularly when several different constraints are present (Araus et al., 2003a; Bassi and Sanchez-Garcia, 2017; Dodig et al., 2012; Maccaferri et al., 2011; Marti and Slafer, 2014; Parent et al., 2017; Slafer et al., 2014; Tardieu and Tuberosa, 2010; Zaim et al. 2017).

To develop durum genotypes with improved yield and adaptability to more frequent incidence of drought and heat stress and/or altered rainfall distribution (Habash et al., 2009; Tadesse et al., 2016), enhancing the genetic background through targeted introgressions is a powerful strategy, given that cultivated germplasm represents only a very small fraction of the variability present in nature (Royo et al., 2009 and references therein; Zaim et al. 2017). Wheat-alien introgression experiments conducted in the past proved to be a valid approach to harness the genetic diversity of alien, mostly wild, segments of wheat-related gene pools (Ceoloni et al., 2014b, 2017a; Dempewolf et al., 2017; Prohens et al., 2017; Zhang et al., 2017). To achieve this, addition/substitution/translocation/recombinant lines, harbouring parts of alien genomes, can be used to facilitate the introduction of desired genes into stable and adapted genotypes (e.g. Ceoloni et al., 2015). Targeted and precise exploitation of useful genes from these materials is readily possible through efficient sexual means, foremost the cytogenetic methodologies of “chromosome engineering” (Ceoloni et al., 2005, 2014a, 2014b). This approach, integrated with continuously developing techniques of genome and chromosome analysis (e.g. marker-assisted selection, association mapping, next generation sequencing, *in situ* hybridization), represents a unique platform for creation of novel and breeder-friendly genetic stocks.

The use of wild relatives for yield improvement in wheat has so far been sporadic, as their productivity is poor, and conspicuous effects on wheat yield rarely observed (Ceoloni et al., 2015; Dempewolf et al., 2017; Zhang et al., 2017). Noteworthy examples regard mostly the hexaploid bread wheat, more widely cultivated, and benefiting from a higher ploidy level with respect to durum wheat, hence a higher buffering ability toward chromosome manipulations (reviewed in Ceoloni et al., 2014a, 2015; Mondal et al., 2016). One of the most notable and documented cases of alien introgression with positive effects on wheat yield, is the transfer of a portion from the group 7 chromosome arm (= 7AgL or 7el_1_L) of the decaploid perennial species *Thinopyrum ponticum* (Popd.) Barkworth & D. R. Dewey (2n = 10x = 70, genomes E^e^E^e^E^x^StSt, see Ceoloni et al., 2014b) onto the 7DL and 7AL arm of bread and durum wheat, respectively. In bread wheat, the sizeable 7AgL translocation named T4 (~70% of the recipient 7DL arm, harbouring *Lr19+Sr25+Yp* genes) led to increased grain yield, biomass and grain number (10-35%) across a number of non-moisture stress environments, and in different backgrounds of CIMMYT germplasm (Monneveux et al., 2003; Reynolds et al., 2001; Singh et al., 1998; Tripathi et al., 2005; Miralles et al., 2007). Under water stress, however, yields for CIMMYT T4 derivatives turned out to be lower than control lines (Monneveux et al., 2003; Singh et al., 1998), as was the case for T4 derivatives developed in Australian adapted genetic backgrounds, when tested in high- and low-yielding environments (Rosewarne et al., 2015).

In durum wheat, three fractions of the same 7AgL chromatin, spanning 23%, 28% and 40% of the recipient 7AL arm of cv. Simeto, and all containing the *Lr19+Sr25+Yp* genes (Ceoloni et al., 2005), were separately introgressed into near-isogenic recombinant lines (NIRLs), and observed across four years in one rain-fed locality of Central Italy (Kuzmanović et al., 2014; 2016). The range of increases in grain yield, biomass and grain number was 3-39%, depending on the recombinant type, season and experimental procedure (spaced plants in Kuzmanović et al., 2014, plot trials in Kuzmanović et al., 2016). In addition, and in contrast to the bread wheat (T4) studies, characterization of the three durum wheat-*Th. ponticum* recombinants comprised more traits, including detailed phenology, spike fertility and flag leaf attributes, and revealed 19 enhanced traits in association with the presence of specific 7AgL portions. This enabled a structural-functional dissection of the 7AgL chromatin incorporated onto the durum 7AL, with consequent assignment of yield-contributing genes (previously associated to the entire T4 segment) to defined 7AgL sub-regions (Kuzmanović et al., 2014, 2016). The increase of several yield-related traits was recorded in each of the three durum wheat-*Th. ponticum* NIRLs. However, the one carrying the 28%-long 7AgL segment was identified as the best performing line, with a high number of yield-related traits (tiller/spike number, flag leaf dimensions and chlorophyll content, grain yield, biomass, duration of stem elongation phase) being evidently enhanced by genetic factor(s) located within the 23-28% chromosomal stretch of its 7AgL segment.

With no information from other environments on the expression of 7AgL and its effects on yield in durum wheat, the aim of the present work was to assess the yield performance of the same three durum wheat-*Th. ponticum* recombinants across an array of rain-fed environments located in three different countries, and to evaluate possible environment/segment-specific associations with final yield and individual yield contributing traits, in view of using these recombinants across environments or in site-directed breeding programs.

## Materials and methods

### Plant materials

Three durum wheat-*Th. ponticum* NIRLs, named R5-2-10, R112-4 and R23-1 (hereafter referred to as R5, R112 and R23, respectively), developed in the background of cv. Simeto by repeated backcrossing (BC) (Ceoloni et al., 2005), were used across four years and three countries. Simeto (pedigree: selection from Capeiti 8 x Valnova) is a variety released in 1988, well adapted to the Italian growing conditions. The NIRLs have portions of *Th. ponticum* 7AgL chromosome arm replacing 23%, 28% and 40% of their distal 7AL arm, respectively, and all three lines include the *Lr19+Yp+Sr25* genes in the sub-telomeric region. Similarly to the plant material described in Kuzmanović et al. (2016), each of the genotypes analysed, corresponding here to BC_5_F_5-9_ (R5 and R112) and BC_4_F_5-9_ (R23) progenies, was represented by either being a homozygous carrier (“+”) or non-carrier (“−”) of the given 7AgL segment. Each “+” and “−“ NIRL included two families originating from sister lines.

### Field experiments

A total of nine rain-fed field trials were carried out over four years and four locations where durum is typically cultivated (two in Italy, one in Morocco and in one Australia) and used for the multi-environment yield assessment. Details on all trials are reported in Table 1. Years and locations were combined and hereafter referred to as environments, with a specific acronym assigned in Table 1. Two of the nine trials have been described previously in Kuzmanović et al. (2016) (VT12 and VT13), from where a subset of traits was considered in the present analysis. In ten environments, all three NIRLs with respective controls were used, while in AUS14, only R5 and R112 were analysed. Sowing densities applied were those commonly used in each of the experimental locations (Table 1). In all field experiments, complete randomized block designs with three replicates for each sister line was used, resulting in a total of 24 plots in AUS14 and 36 plots in the other eight environments (2 per each sister line, i.e. 6 per each +/− NIRL). Meteorological data during the growing seasons for daily temperatures (minimum, mean and maximum) and rainfall (Table 2) were retrieved from meteorological stations at experimental sites, except for MOR14, for which the data were downloaded from NASA’s (National Aeronautics and Space Administration, USA) site for Prediction of Worldwide Energy Resource (http://power.larc.nasa.gov). All trials were managed according to standard local practices including fertilization, weed, pest and disease control, in order to avoid, in particular, leaf rust spreading on *Lr19* non-carrier plants (- NIRLs), hence to eliminate the indirect yield-contributing factor of *Lr19*-carriers (+ NIRLs).

**Table 1.** Description of locations and field experiments analysed in this study (NIRL, Near Isogenic Recombinant Line).

| Environment acronym | Location | Latitude | Longitude | Altitude (m) | Season | Total rainfall (mm) | Mean temperature at heading (C°) | NIRLs tested | Sowing date | Crop cycle length (days) | Sowing density (seed/m<sup>2</sup>) | Plot size (m<sup>2</sup>) |
| --- | --- | --- | --- | --- | --- | --- | --- | --- | --- | --- | --- | --- |
| VT12 | Experimental farm of the University of Tuscia, Viterbo (Central Italy) | 42° 25' N | 12° 4' E | 301 | 2011/12 | 248 | 13.6 | R5, R112, R23 | 17/11/2011 | 226 | 350 | 2.3 |
| VT13 | " | " | " | " | 2012/13 | 534 | 17.2 | " | 20/12/2012 | 192 | 350 | 2.3 |
| VT14 | " | " | " | " | 2013/14 | 676 | 12.7 | " | 29/11/2013 | 213 | 350 | 4.5 |
| VT15 | " | " | " | " | 2014/15 | 337 | 16.5 | " | 15/12/2014 | 197 | 350 | 4.5 |
| BO14 | Experimental farm of the University of Bologna, Cadriano (North Italy) | 44° 33' N | 11° 24' E | 33 | 2013/14 | 598 | 13.1 | " | 07/11/2013 | 236 | 350 | 4.2 |
| MOR14 | Marchouch (Morocco) | 33° 36' N | -6° 43' W | 440 | 2013/14 | 229 | 13.0 | " | 03/12/2013 | 189 | 300 | 6 |
| MOR15 | " | " | " | " | 2014/15 | 349 | 14.1 | " | 18/11/2014 | 204 | 300 | 6 |
| AUS13 | Waite campus, University of Adelaide, Urrbrae (South Australia) | 34° 58' S | 138° 38' E | 48 | 2013 | 305 | 16.9 | " | 19/06/2013 | 179 | 150 | 1.6 |
| AUS14 | " | " | " | " | 2014 | 296 | 14.3 | R5, R112 | 15/05/2014 | 193 | 150 | 5 |

**Table 2.**
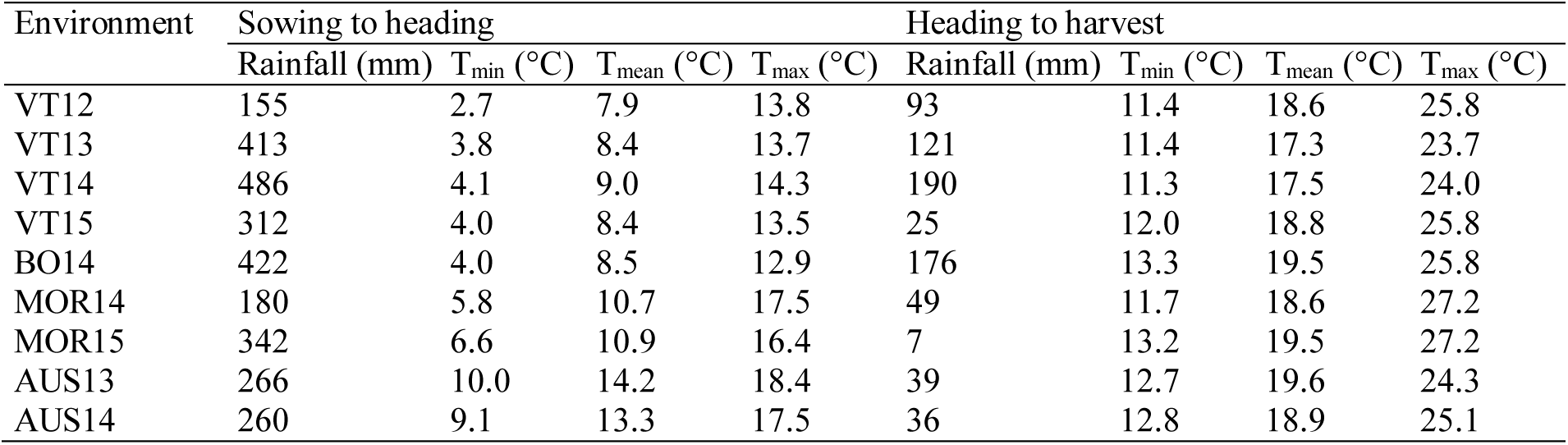
Weather conditions for growing seasons in nine environments analysed, as retrieved from meteorological stations at experimental sites, or in the case of MOR14 only, downloaded from NASA’s site for Prediction of Worldwide Energy Resource (http://power.larc.nasa.gov). Environment acronyms are as described in Table 1.

| Environment | Sowing to heading |  |  |  | Heading to harvest |  |  |  |
| --- | --- | --- | --- | --- | --- | --- | --- | --- |
|  | Rainfall (mm) | T<sub>min</sub> (°C) | T<sub>mean</sub> (°C) | T<sub>max</sub> (°C) | Rainfall (mm) | T<sub>min</sub> (°C) | T<sub>mean</sub> (°C) | T<sub>max</sub> (°C) |
| VT12 | 155 | 2.7 | 7.9 | 13.8 | 93 | 11.4 | 18.6 | 25.8 |
| VT13 | 413 | 3.8 | 8.4 | 13.7 | 121 | 11.4 | 17.3 | 23.7 |
| VT14 | 486 | 4.1 | 9.0 | 14.3 | 190 | 11.3 | 17.5 | 24.0 |
| VT15 | 312 | 4.0 | 8.4 | 13.5 | 25 | 12.0 | 18.8 | 25.8 |
| BO14 | 422 | 4.0 | 8.5 | 12.9 | 176 | 13.3 | 19.5 | 25.8 |
| MOR14 | 180 | 5.8 | 10.7 | 17.5 | 49 | 11.7 | 18.6 | 27.2 |
| MOR15 | 342 | 6.6 | 10.9 | 16.4 | 7 | 13.2 | 19.5 | 27.2 |
| AUS13 | 266 | 10.0 | 14.2 | 18.4 | 39 | 12.7 | 19.6 | 24.3 |
| AUS14 | 260 | 9.1 | 13.3 | 17.5 | 36 | 12.8 | 18.9 | 25.1 |

### Measurements of yield-related traits

A list of traits and details on environments in which the materials were analysed, as well as on the sample type and number for each replicated plot, are reported in Table 3. Measurements of all traits were performed as described in Kuzmanović et al. (2016) with some modifications. HD was considered as number of days from sowing to heading. Samples of culms and flag leaves were randomly chosen within each plot. All dry weights of culms, spikes, chaff and total aboveground biomass were recorded after 48 h oven drying at 65°C. SIA was calculated as SDWA/TDWA ratio, SFI as the ratio between GNS and CHAFF at maturity (Isidro et al., 2011). SFI is a spike fertility parameter positively correlated with fruiting efficiency (Abbate et al., 2013; Martino et al., 2015), calculated as the ratio between GNS and SDWA (e.g., Slafer et al., 2015; Terrile et al., 2017). FLA was determined using the formula FLW × FLL × 0.75 (Dodig et al., 2010). Chlorophyll content was measured by using a hand-held meter SPAD 502 (Konica-Minolta, Japan) at medium milk (Zadoks 75; Zadoks et al., 1974), late milk (Zadoks 77) and very late milk (Zadoks 79) developmental stages. HI was determined as GYM2/BM2 in MOR14, MOR15, AUS13 and AUS14, where GYM2 and BM2 were obtained directly from the total harvested area. In other environments, HI was determined on 25-culm samples plot^−1^, and at harvest used for BM2 estimation, once the total plot area was trashed and weighed (=GYM2/HI). TGW was obtained from weighing two 100-seed samples plot^−1^ and then used to evaluate GNM2 (=GYM2 × 1000/TGW).

**Table 3.**
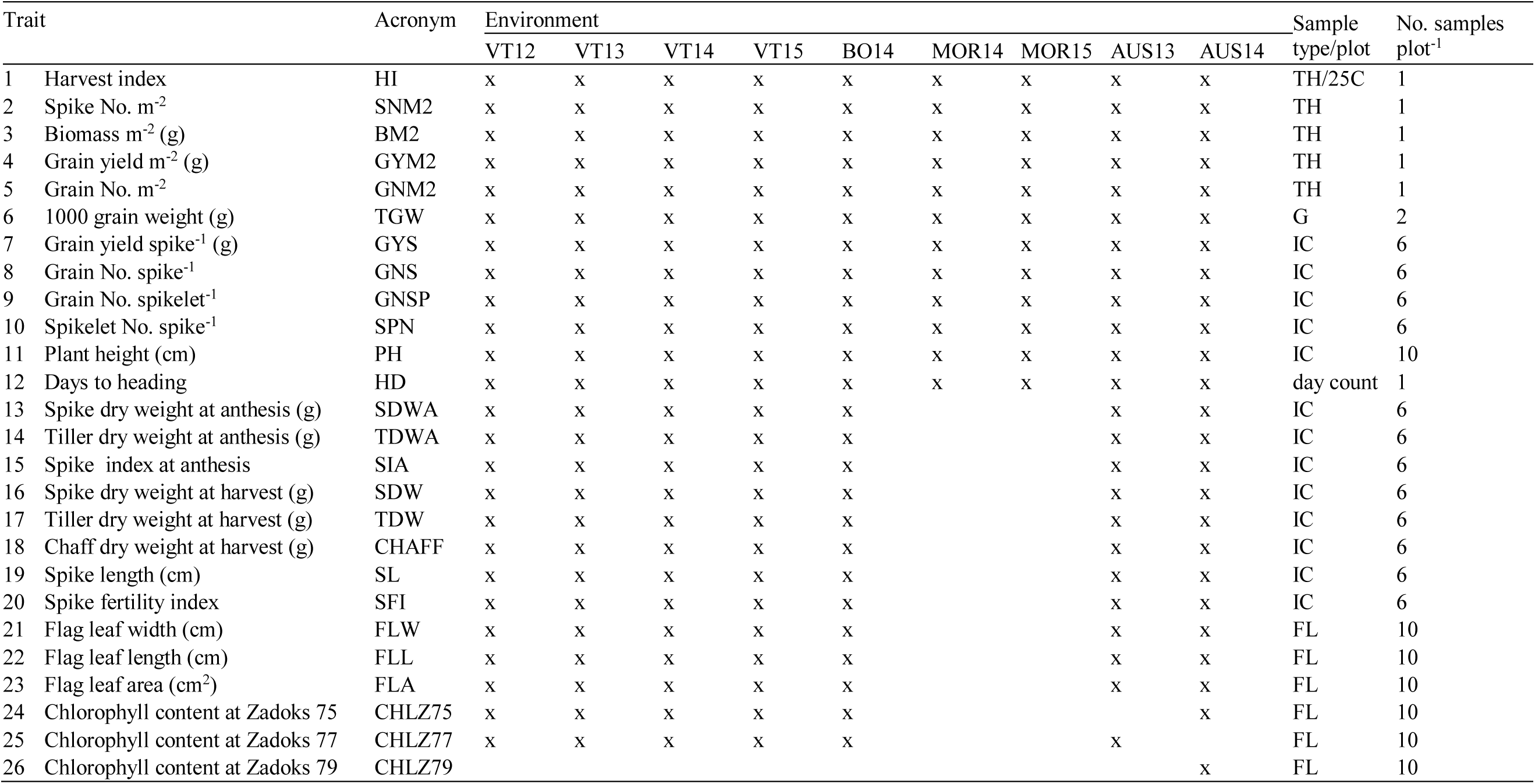
Traits and sample details assessed across nine environments analysed in this study [TH, total plot harvest; 25C, 25 culms; G, grain sample (1 or 2); IC, individual culms (5-10); FL, individual flag leaves (5-10)]. Environment acronyms are as described in Table 1.

### Statistical analyses

All analyses were performed by SYSTAT12 software (Systat Software Incorporated, San Jose, CA, USA). To investigate the effects of genetic and environmental factors and interactions between them on recorded traits, an analysis of variance (ANOVA) was performed, applying to datasets a general linear model (GLM) as a mixed effect model. Three such models were employed (= GLM1-3), depending on the data subset taken into consideration, as not all of the 25 traits and not all of the three NIRLs were analysed in all nine environments. In GLM1 and GLM2, datasets comprised environments where all three NIRLs were tested, while in GLM3, only the AUS14 dataset was analysed. GLM1 included traits No. 1-12 listed in Table 3, GLM2 traits No. 1-25, while GLM3 comprised traits No. 1-24 and 26. GLM1 was applied for the analysis of the overall effect of presence/absence of alien segments on main yield-related traits across environments while GLM2 and GLM3 were applied for the analysis of individual 7AgL segment effects. Each variable (i.e. trait measured) was entered as a ‘dependent’ factor against ‘independent’ factors. The latter were: genotype background (G), i.e. background genetic information from the recurrent variety, environment (E), presence/absence of the 7AgL segment [7AgL alone in GLM1 or 7AgL(G), i.e. nested in the background, in other GLMs], and replicate [R alone in GLM3 and environment-nested, R(E), in other GLMs]. The latter factor was used in the models as the error. First order [E × G; R(E); E × 7AgL; 7AgL(G)], and second order [E × 7AgL(G)] interactions between the above factors were analysed as well. In all analyses three levels of significance were considered, corresponding to *P* < 0.05, *P* < 0.01 and *P* < 0.001. When significant factors and/or interactions between them (*F* values) were observed, a pairwise analysis was carried out by the Tukey Honestly-Significant-Difference test at the 0.95 confidence level.

A correlation matrix was built for a subset of traits recorded in all environments. Each pair of variables was correlated by calculating Pearson’s correlation coefficients (*r* value), while the significance levels were obtained using the Bonferroni method. Simple linear regression analysis was carried out by applying the least squares method for data fitting a 0.95 confidence level.

In order to analyse any genotype-specific response for grain yield (GYM2) across environments, the environmental index was calculated (= average value of all participating genotypes) in each environment and combined with the environmental mean of each genotype in a simple linear regression (b coefficient statistics). GYM2 values were transformed in logarithmic to attain linearity and homogeneity of errors (Finlay and Wilkinson 1963) and to observe the relative (“intrinsic”) variability of interest (Becker and Leon, 1988).

## Results

### Environments

The field trials described here cover a wide range of environmental conditions, as usually observed in areas of durum wheat cultivation. Overall, environments were comparable for mean temperatures (Table 2), with a little more variation between them in the period from sowing to heading (6.3°C) than from heading to maturity (2.3°C). By contrast, the environments were considerably different for the rainfall amount received (Table 1) and its distribution during the growing season (Table 2). Large differences in rainfall input were recorded particularly from heading to maturity, being in the range from 7 to 190 mm. The two seasons in South Australia were virtually identical for the precipitation received, nearly approaching the average values of the agricultural southeast areas of the State, while being the warmest years on record (since 1910), particularly through the second half of the season (www.bom.gov.au). The two trials in Morocco were also characterized by very warm seasons, with average daily maximum temperatures from May until harvest, often exceeding the site’s average daily maximum values (22-26°C) by up to 16°C (www.weatherspark.com). Although less abundant, precipitation events were better distributed in MOR14 than in MOR15 (Table 2). As for the Italian trial sites, season 2012 was more typical and very favourable for durum wheat cultivation when compared to the 2013 to 2015 seasons. The latter three seasons were characterized by exceptional (and numerous) precipitation events and an overall increase in temperature during the entire crop cycle (www.informatoreagrario.it). VT12 had rainfall distribution and temperature trends that were conducive for good crop growth during the entire life cycle (Table 2; Kuzmanović et al., 2016). Still, in VT12, drought stress was present with precipitation amounts significantly below the site’s mean, and higher maximum temperatures during the second part of the growth cycle (www.informatoreagrario.it). Conversely, seasons 2013 to 2015 in Italy had unusually rainy and mild winters, with a full soil moisture profile. Particularly heavy and prolonged rain periods prior to sowing and during the grain filling period occurred in VT13, VT14 and BO14, while VT15 resulted in very dry and hot conditions from heading onwards with respect to the former three environments and to the site’s mean values.

### Yield response across environments

According to the observed highly significant R^2^ values of linear regression (Fig. 1a), all NIRLs (both + and −) positively responded to better thermo-pluviometric patterns and higher environmental indices across environments. The b coefficient values around 1 for R5 and R112 genotypes were indicative of an average yield stability (Finlay and Wilkinson, 1963), and the consistently higher grain yield with respect to the site’s mean indicated their general adaptability. In contrast, the R23 NIRL pair, despite b coefficient values around 1 (i.e. average stability indicator), had poorer grain yield than the site’s mean in all environments, probably due to the lower adaptability of the genetic background of its representative families, less isogenic compared to the recurrent cv. Simeto parent than the two other NIRLs (see § 2.1).

**Fig. 1.**
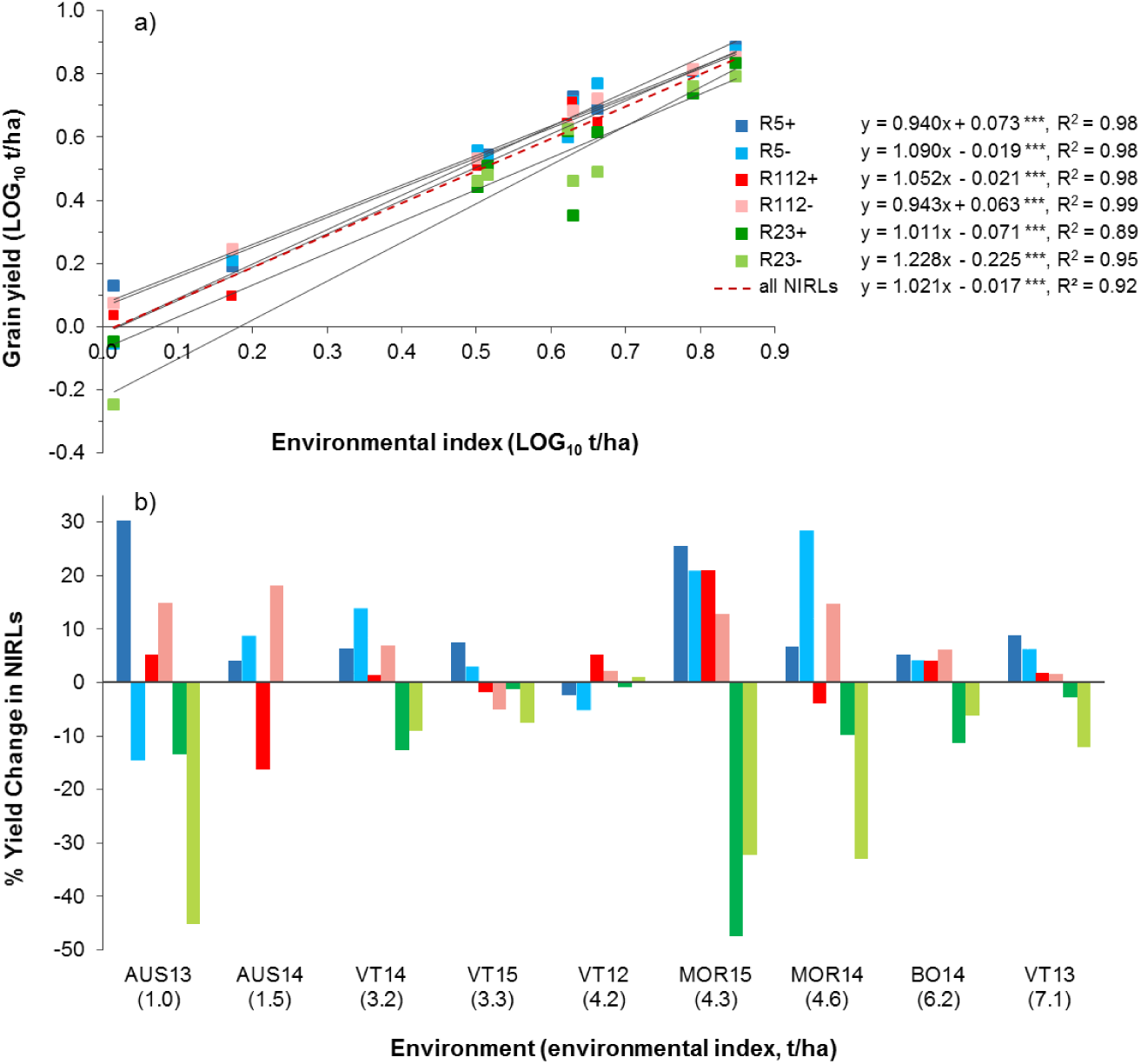
Regression lines showing the relationship of individual grain yields of the six NIRLs and nine environments analysed (a), and the percentage yield change of the six NIRLs with respect to the site’s mean (b). Environment acronyms are as described in Table 1.

With an average grain yield across environments between 1.03 and 7.05 t/ha (Fig. 1b), environments were arbitrarily classified as: low-yielding (grain yield lower than 2 t/ha), medium-yielding (grain yield between 2 and 5 t/ha), and high-yielding (grain yield higher than 5 t/ha). The productivity of R5 and R112 NIRLs was typically higher than the environmental means, as shown from their yield gains ranging 3-30% and 2-27%, respectively. Yield of R23 NIRLs (both “+” and “−”) was always under the site’s mean (−1 to −47%). Yet, in four out of eight environments (all but AUS14, Table 1), the presence of the R23 7AgL segment evidently had a positive effect, reducing the background-dependent yield disadvantage (Fig. 1b). The two more productive 7AgL-carrier lines (R5+ and R112+), displayed yield gains with respect to their control NIRLs in 5 and 3, respectively, out of the nine environments. The highest gain of R5+ (and of all 7AgL+ lines) was observed in low-yielding AUS13 (+30%), a similar value to that recorded in medium-yielding MOR15 (+26%). In the same MOR15, R112+ showed the highest gain (+21%). By contrast, a notable yield penalty was observed for R112+ in low-yielding AUS14 (−18%), and for R23+ in medium-yielding MOR15 (−47%).

### 7AgL-associated effects on yield and yield-related traits

Across all eight environments where all three NIRLs were tested (all but AUS14, Table 1), when data from the three “+” recombinant or three “−“ control lines were pooled, yield parameters turned out to be increased to a variable extent in 7AgL+ vs. 7AgL− genotypes (Table 4 and Supplementary Table 1). The most significant positive 7AgL effects were observed for SNM2 (+13%), BM2 (+9%), GNM2 (+11%), and GNSP (+8%) accompanied though by a significant decrease in TGW and GYS. However, HI of 7AgL+ lines remained unchanged (Table 4). Although the environment factor alone (E, Supplementary Table 1) was highly significant for all traits, confirming that the environments tested were different, the association between E and 7AgL turned out to be not significant for any of the traits (as observed from the Tukey test for E × 7AgL, data not shown). ANOVA performed for each of the nine environments separately, in view of highlighting the 7AgL effects in the various contrasting conditions, revealed specific associations between positive 7AgL-linked effects and sites (Table 4). SNM2, GNM2 and GNS were significantly increased in 7AgL+ vs. 7AgL− NIRLs in several environments, particularly in the lowest-(AUS13) and highest-yielding (BO14, VT13), as well as in the medium-yielding VT12. AUS13 turned out to be the only environment in which GYM2 together with BM2 were significantly increased (+18% and +14%, respectively) in 7AgL+ vs. 7AgL− NIRLs. This may indicate potential for adaptation to stressed conditions conferred by the 7AgL introgressions.

**Table 4.** Main yield-related traits unit area^−1^ and spike^−1^ of 7AgL-carrier vs. 7AgL-non carrier durum wheat-*Th. ponticum* NIRLs across environments (HI, harvest index; SNM2, spike number m^−2^; BM2, biomass m^−2^; GYM2, grain yield m^−2^; GNM2, grain number m^−2^; TGW, thousand grain weight; GYS, grain yield spike^−1^; GNS, grain number spike^−1^; GNSP, grain No. spikelet^−1^; SPN, spikelet No. spike^−1^; PH, plant height; HD, heading date). In eight out of the nine environments, all three recombinants and their respective controls were analysed (ALL); in AUS14, only R5 and R112 NIRLs were included. Positive and negative 7AgL effects are highlighted in *green* and *red*, respectively. \**P* <0.05, \*\**P*<0.01, \*\*\**P* <0.001. Environments are ordered from the lowest- to the highest-yielding one, and their acronyms are as described in Table 1.

| Trait | NIRLs | Environment |  |  |  |  |  |  |  |  |  |
| --- | --- | --- | --- | --- | --- | --- | --- | --- | --- | --- | --- |
|  |  | ALL | AUS13 | AUS14 | VT14 | VT15 | VT12 | MOR15 | MOR14 | BO14 | VT13 |
| HI | 7AgL+ | 0.359 | 0.234 | 0.375 | 0.424 | 0.453 | 0.406 | 0.262 | 0.297 | 0.442 | 0.482 |
|  | 7AgL- | 0.360 | 0.224 | 0.420 | 0.410 | 0.466 | 0.412 | 0.289 | 0.306 | 0.447 | 0.483 |
|  | P-value | 0.516 | 0.420 | 0.056 | 0.351 | 0.578 | 0.484 | 0.533 | 0.666 | 0.790 | 0.943 |
|  | 7AgL effect (%) | 0 | 4 | -11 | 3 | -3 | -1 | -9 | -3 | -1 | 0 |
| SNM2 | 7AgL+ | 262.6 | 121.3 | 54.9 | 188.6 | 179.8 | 284.9 | 371.8 | 338.5 | 372.4 | 370.5 |
|  | 7AgL- | 233.3 | 91.6 | 52.9 | 187.0 | 164.9 | 259.2 | 356.9 | 354.0 | 343.0 | 342.2 |
|  | P-value | 0.009** | 0.000*** | 0.865 | 0.880 | 0.265 | 0.036* | 0.684 | 0.643 | 0.041* | 0.028* |
|  | 7AgL effect (%) | 13 | 32 | 4 | 1 | 9 | 10 | 4 | -4 | 9 | 8 |
| BM2 | 7AgL+ | 1064.3 | 474.1 | 376.3 | 726.3 | 739.1 | 1041.8 | 1681.3 | 1570.7 | 1397.0 | 1512.2 |
|  | 7AgL- | 979.6 | 416.2 | 402.5 | 794.3 | 679.4 | 1012.5 | 1487.5 | 1543.0 | 1399.6 | 1443.2 |
|  | P-value | 0.018* | 0.001** | 0.583 | 0.218 | 0.254 | 0.471 | 0.065 | 0.636 | 0.973 | 0.120 |
|  | 7AgL effect (%) | 9 | 14 | -7 | -9 | 9 | 3 | 13 | 2 | 0 | 5 |
| GYM2 | 7AgL+ | 384.8 | 110.9 | 136.8 | 311.1 | 332.6 | 421.5 | 424.9 | 467.0 | 612.2 | 721.2 |
|  | 7AgL- | 372.6 | 94.3 | 169.0 | 330.2 | 316.8 | 416.4 | 428.3 | 475.4 | 625.4 | 688.6 |
|  | P-value | 0.782 | 0.029* | 0.115 | 0.433 | 0.304 | 0.752 | 0.959 | 0.844 | 0.685 | 0.281 |
|  | 7AgL effect (%) | 3 | 18 | -19 | -6 | 5 | 1 | -1 | -2 | -2 | 5 |
| GNM2 | 7AgL+ | 7871.2 | 1132.5 | 3058.1 | 5681.6 | 5703.0 | 15589.9 | 9349.4 | 10297.3 | 10234.4 | 12297.9 |
|  | 7AgL- | 7100.5 | 813.1 | 3323.2 | 5544.7 | 5101.9 | 13635.0 | 8488.2 | 9188.1 | 9526.1 | 10425.8 |
|  | P-value | 0.000*** | 0.000*** | 0.510 | 0.903 | 0.172 | 0.038* | 0.353 | 0.110 | 0.172 | 0.001** |
|  | 7AgL effect (%) | 11 | 39 | -8 | 2 | 12 | 14 | 10 | 12 | 7 | 18 |
| TGW | 7AgL+ | 50.8 | 34.8 | 46.6 | 55.4 | 59.2 | 60.5 | 46.4 | 45.6 | 60.1 | 60.3 |
|  | 7AgL- | 55.3 | 38.1 | 51.0 | 59.9 | 62.3 | 66.8 | 49.7 | 51.1 | 65.9 | 66.3 |
|  | P-value | 0.000*** | 0.006** | 0.148 | 0.097 | 0.417 | 0.032* | 0.274 | 0.048* | 0.021* | 0.005** |
|  | 7AgL effect (%) | -8 | -9 | -9 | -8 | -5 | -9 | -7 | -11 | -9 | -9 |
| GYS | 7AgL+ | 2.3 | 1.0 | 2.7 | 3.1 | 2.9 | 2.6 | 3.1 | 2.2 | 2.3 | 2.9 |
|  | 7AgL- | 2.4 | 1.1 | 3.2 | 3.0 | 3.2 | 2.8 | 3.5 | 2.3 | 2.6 | 3.0 |
|  | P-value | 0.000*** | 0.168 | 0.022* | 0.595 | 0.130 | 0.041* | 0.071 | 0.368 | 0.124 | 0.269 |
|  | 7AgL effect (%) | -4 | -8 | -18 | 3 | -9 | -6 | -13 | -6 | -11 | -4 |
| GNS | 7AgL+ | 43.1 | 24.6 | 59.4 | 54.8 | 49.8 | 43.4 | 52.6 | 46.9 | 42.3 | 47.7 |
|  | 7AgL- | 39.7 | 21.8 | 66.3 | 47.7 | 51.3 | 41.4 | 56.6 | 44.9 | 43.9 | 43.7 |
|  | P-value | 0.057 | 0.028* | 0.023* | 0.004** | 0.460 | 0.280 | 0.142 | 0.358 | 0.532 | 0.006** |
|  | 7AgL effect (%) | 9 | 13 | -10 | 15 | -3 | 5 | -7 | 5 | -3 | 9 |
| GNSP | 7AgL+ | 2.4 | 1.2 | 2.4 | 3.0 | 2.9 | 2.5 | 2.9 | 2.4 | 2.2 | 3.0 |
|  | 7AgL- | 2.2 | 1.1 | 2.2 | 2.6 | 2.9 | 2.4 | 2.8 | 2.3 | 2.3 | 2.8 |
|  | P-value | 0.005** | 0.078 | 0.072 | 0.007** | 0.814 | 0.185 | 0.462 | 0.547 | 0.594 | 0.021* |
|  | 7AgL effect (%) | 8 | 11 | 7 | 13 | -1 | 4 | 4 | 3 | -3 | 7 |
| SPN | 7AgL+ | 18.7 | 20.9 | 18.8 | 18.5 | 17.2 | 17.2 | 18.4 | 19.7 | 19.2 | 15.9 |
|  | 7AgL- | 18.7 | 20.9 | 18.9 | 18.1 | 17.7 | 17.1 | 20.7 | 19.7 | 19.4 | 15.7 |
|  | P-value | 0.145 | 0.910 | 0.904 | 0.512 | 0.233 | 0.703 | 0.044* | 0.965 | 0.613 | 0.485 |
|  | 7AgL effect (%) | 0 | 0 | 0 | 2 | -3 | 1 | -11 | 0 | -1 | 2 |
| PH | 7AgL+ | 82.4 | 74.7 | 82.6 | 91.8 | 75.1 | 87.1 | 85.0 | 84.7 | 86.2 | 82.4 |
|  | 7AgL- | 81.4 | 72.3 | 81.7 | 90.9 | 76.2 | 87.0 | 89.4 | 83.9 | 86.4 | 81.3 |
|  | P-value | 0.965 | 0.214 | 0.490 | 0.862 | 0.777 | 0.984 | 0.357 | 0.776 | 0.960 | 0.773 |
|  | 7AgL effect (%) | 1 | 3 | 1 | 1 | -1 | 0 | -5 | 1 | 0 | 1 |
| HD | 7AgL+ | 130.0 | 99.4 | 129.5 | 142.2 | 139.3 | 153.6 | 131.7 | 105.8 | 166.4 | 132.0 |
|  | 7AgL- | 127.0 | 95.3 | 126.6 | 141.0 | 138.7 | 153.2 | 130.4 | 105.7 | 165.6 | 130.6 |
|  | P-value | 0.011* | 0.000*** | 0.295 | 0.499 | 0.334 | 0.575 | 0.609 | 0.904 | 0.562 | 0.011* |
|  | 7AgL effect (%) | 2 | 4 | 2 | 1 | 0 | 0 | 1 | 0 | 1 | 1 |

The more extensive and detailed GLM2 model (Supplementary Table 2), used to examine individual effects of each of the three 7AgL segments (Table 5), showed that E alone was highly significant for all traits, but also the majority of interactions including it. Across all environments, the influence of the individual 7AgL segments on the various traits [7AgL(G), Supplementary Table 2] was different, as revealed by analysis of the three “+” vs. “−” NIRL pairs. Taken the eight environments where all three recombinants were analysed (all but AUS14, Table 1) as a whole, the R5+ recombinant did not show any significant trait change due to its 7AgL segment (Table 5), although in the AUS13 environment alone several significant differences emerged (see § 3.3.1). Also the comparison between R112+ and its R112− control across all environments showed only minor differences for HI (−3%), TGW (−4%), HD (+4%) and FLL (−9%) (Table 5).

**Table 5.** Mean values and standard errors (SE) of yield-related traits of the three durum wheat-*Th. ponticum* NIRLs (+, 7AgL carriers; −, 7AgL non-carriers) across eight environments where all three recombinants and their respective controls were analysed (HI, harvest index; SNM2, spike number m^−2^; BM2, biomass m^−2^; GYM2, grain yield m^−2^; GNM2, grain number m^−2^; TGW, thousand grain weight; GYS, grain yield spike^−1^; GNS, grain number spike^−1^; GNSP, grain number spikelet^−1^; SPN, spikelet number spike^−1^; PH, plant height; HD, heading date; SDWA, spike dry weight at anthesis; TDWA, tiller dry weight at anthesis; SIA, spike index at anthesis; SDW, spike dry weight at maturity; TDW, tiller dry weight at maturity; CHAFF, chaff dry weight; SL, spike length; SFI, spike fertility index; FLW, flag leaf width; FLL, flag leaf length; FLA, flag leaf area; CHLZ75, flag leaf chlorophyll content at Zadoks 75; CHLZ77, flag leaf chlorophyll content at Zadoks 77). Letters in each row correspond to the ranking of the Tukey test at *P* < 0.01 (capital) and *P* < 0.05 (lower case) levels.

| Trait | R5+ |  |  | R5- |  |  | R112+ |  |  | R112- |  |  | R23+ |  |  | R23- |  |  |
| --- | --- | --- | --- | --- | --- | --- | --- | --- | --- | --- | --- | --- | --- | --- | --- | --- | --- | --- |
|  | Mean | SE |  | Mean | SE |  | Mean | SE |  | Mean | SE |  | Mean | SE |  | Mean | SE |  |
| HI | 0.39 | 0.01 | a | 0.38 | 0.02 | a | 0.37 | 0.01 | b | 0.38 | 0.02 | a | 0.31 | 0.02 | c | 0.32 | 0.02 | c |
| SNM2 | 251.1 | 16.0 | AB | 228.8 | 19.6 | AB | 263.0 | 15.9 | AB | 230.4 | 18.9 | AB | 273.5 | 15.8 | A | 241.8 | 18.6 | B |
| BM2 | 1041.3 | 62.1 | ns | 932.1 | 76.4 | ns | 1067.1 | 73.7 | ns | 938.5 | 73.8 | ns | 1084.1 | 65.8 | ns | 1079.1 | 76.9 | ns |
| GYM2 | 418.3 | 29.4 | ns | 382.6 | 39.7 | ns | 396.0 | 28.7 | ns | 373.0 | 35.9 | ns | 340.9 | 27.1 | ns | 360.5 | 34.3 | ns |
| GNM2 | 7376.0 | 591.9 | B | 6929.3 | 775.9 | B | 7660.7 | 656.7 | B | 6956.7 | 807.2 | B | 8576.8 | 803.4 | A | 7458.8 | 849.2 | C |
| TGW | 57.5 | 1.5 | A | 56.0 | 2.1 | A | 53.2 | 1.7 | B | 55.6 | 2.0 | A | 41.6 | 1.1 | C | 54.2 | 2.0 | B |
| GYS | 2.6 | 0.1 | A | 2.3 | 0.2 | A | 2.4 | 0.1 | A | 2.4 | 0.2 | A | 2.0 | 0.1 | B | 2.5 | 0.2 | A |
| GNS | 42.5 | 1.4 | ab | 38.8 | 1.9 | b | 42.5 | 1.6 | ab | 40.1 | 2.0 | ab | 44.1 | 1.9 | a | 40.2 | 2.6 | b |
| GNSP | 2.4 | 0.1 | A | 2.2 | 0.1 | A | 2.3 | 0.1 | A | 2.3 | 0.1 | A | 2.3 | 0.1 | A | 2.1 | 0.1 | B |
| SPN | 18.2 | 0.3 | C | 18.3 | 0.4 | C | 18.6 | 0.3 | BC | 18.1 | 0.3 | C | 19.2 | 0.3 | B | 20.0 | 0.5 | A |
| PH | 77.9 | 0.9 | c | 75.8 | 1.0 | cd | 76.1 | 0.8 | d | 73.2 | 0.9 | d | 93.2 | 1.2 | b | 97.8 | 1.4 | a |
| HD | 127.2 | 3.1 | C | 123.6 | 4.0 | C | 129.2 | 3.2 | B | 123.7 | 3.9 | C | 133.6 | 3.1 | A | 135.0 | 3.7 | A |
| SDWA | 0.76 | 0.03 | AB | 0.79 | 0.04 | AB | 0.74 | 0.03 | AB | 0.78 | 0.03 | A | 0.55 | 0.02 | C | 0.66 | 0.02 | B |
| TDWA | 4.46 | 0.14 | A | 4.82 | 0.20 | A | 4.39 | 0.14 | A | 4.51 | 0.15 | A | 3.83 | 0.14 | B | 4.65 | 0.22 | A |
| SIA | 0.17 | 0.01 | ns | 0.16 | 0.01 | ns | 0.17 | 0.00 | ns | 0.17 | 0.00 | ns | 0.15 | 0.00 | ns | 0.15 | 0.01 | ns |
| SDW | 3.26 | 0.12 | A | 3.17 | 0.12 | A | 3.20 | 0.12 | A | 3.17 | 0.12 | A | 2.57 | 0.11 | B | 3.02 | 0.16 | A |
| TDW | 5.26 | 0.21 | b | 4.99 | 0.23 | b | 5.09 | 0.23 | b | 4.92 | 0.25 | b | 4.75 | 0.20 | c | 6.19 | 0.18 | a |
| CHAFF | 0.87 | 0.04 | ab | 1.02 | 0.10 | ab | 0.94 | 0.05 | a | 0.93 | 0.05 | ab | 0.74 | 0.04 | b | 0.90 | 0.04 | a |
| SL | 6.5 | 0.1 | ns | 6.7 | 0.2 | ns | 6.4 | 0.1 | ns | 6.3 | 0.1 | ns | 7.2 | 0.1 | ns | 7.3 | 0.2 | ns |
| SFI | 50.5 | 2.7 | B | 48.1 | 4.3 | B | 51.9 | 3.1 | B | 48.3 | 4.0 | B | 68.5 | 4.6 | A | 46.9 | 3.9 | B |
| FLW | 1.8 | 0.03 | A | 1.7 | 0.04 | AB | 1.9 | 0.03 | AB | 1.8 | 0.03 | B | 1.7 | 0.02 | B | 1.7 | 0.04 | C |
| FLL | 20.3 | 0.59 | ab | 21.7 | 0.90 | ab | 19.9 | 0.59 | bc | 21.8 | 0.90 | a | 18.8 | 0.57 | c | 20.9 | 0.79 | a |
| FLA | 31.1 | 1.67 | A | 30.0 | 1.80 | A | 31.1 | 1.70 | A | 32.0 | 1.80 | A | 26.3 | 1.39 | B | 30.0 | 2.11 | A |
| CHLZ75 | 49.8 | 1.6 | ns | 53.1 | 1.6 | ns | 49.5 | 1.5 | ns | 51.0 | 2.0 | ns | 41.8 | 1.3 | ns | 45.1 | 1.4 | ns |
| CHLZ77 | 42.8 | 1.2 | A | 43.1 | 1.5 | A | 44.4 | 1.3 | A | 42.6 | 1.5 | AB | 39.3 | 1.2 | B | 43.4 | 1.4 | AB |

On the other hand, R23+ was the recombinant for which the highest number of significant differences with respect to its R23− control were observed irrespective of the E factor (Table 5). While a number of yield-related parameters were depressed by the presence of the 40%-long 7AgL segment [TGW, GYS, SPN, PH, dry weight at anthesis and maturity (SDWA, TDWA, SDW, TDW, CHAFF), flag leaf dimensions (FLL, FLA)], the same 7AgL portion was clearly associated with significant increases of several other parameters directly contributing to grain yield, i.e. SNM2 (+13%), GNM2 (+15%), GNS (+10%), GNSP (+10%) and SFI (+46%) (Table 5).

### 7AgL × E interaction

Individual alien segment effects were significant to a variable extent in particular environments, and for specific traits, as revealed by the Tukey test (data not shown) for significant E × 7AgL(G) interactions (GLM2, Supplementary Tables 2 and 3). From this analysis across environments, significant differences between R5+ and R5− control were observed in the low-yielding AUS13 environment. These included increase of HI (+27%) and FLW (+15%), as well as decrease of FLL (−13%) and CHAFF (−38%). Considering the AUS13 dataset alone, additional significant differences in favour of R5+ could be highlighted, namely a 52% increase in grain yield (GYM2), paralleled by significantly higher values of GYS (+30%), GNM2 (+45%), GNS (+21%), SFI (+77%) and BM2 (+21%) (see Supplementary Table 3). As for the R112+ recombinant, the multi-environment dataset showed TGW (−13%), FLL (−15%) and FLW (−17%) to be significantly depressed compared to the R112− control in AUS13, while biomass production. (BM2) to be boosted in MOR15 (+24%) Data from AUS14, in which R5 and R112 NIRLs only were grown, revealed that presence of both 7AgL segments was not particularly advantageous for the main yield traits (traits No. 1-8 from Table 3; Supplementary Table 3). The GLM3 model identified only a few significant differences associated with the presence of the 7AgL segment ([7Ag(G)]; Supplementary Table 4): incremental, for FLW of both recombinants (+37% for R5+ and +25% for R112+), FLA of R5+ (+42%) and HD of R112+ (+6%); detrimental, for GNSP of R112+ (−14%) and for CHL75 of R5+ (−7%) (Supplementary Table 3).

On the other hand, the R23+ NIRL was highly responsive to different environments for several traits. The significant enhancing effect of its 7AgL segment for GNM2 and SFI was particularly pronounced in VT12-VT13 (+30% and +39%, respectively) and MOR14 (+56%) for the former trait, while in VT13-VT14 (+35% and +59%, respectively) and BO14 (+95%) for the latter (Supplementary Table 3). Some of the identified depressing effects of the 40%-long 7AgL segment were significantly associated with certain, even contrasting environments: TGW in VT12-15, BO14 and AUS13 (ranging −18 to −27%); GYS in MOR15 (−43%); dry weight at maturity (SDW, TDW) in BO14 (−27% and −28%, respectively); FLA in BO14 and VT14 (−22% and −33%, respectively). Similarly to the R5 case, analysis of the AUS13 dataset alone showed significant increases of GYM2 (+58%), GNM2 (over 3 fold), SNM2 (+128%), GNS (+49%), GNSP (+67%) and BM2 (+33%) to be also associated to R23+ vs. R23−. as well as significant reduction of TGW (−19%), GYS (−30%), TDWA (−19%) and TDW (−43%) (Supplementary Table 3).

### Correlations of yield traits

Scatter plots of regression analysis (R^2^) for the pairs of main yield traits are shown in Fig. 2, and coefficients of correlation (*r*) are reported in Table 6. For all environments, final grain yield per area (GYM2) was mostly dependent on SNM2, BM2 and GNM2 (R^2^ = 0.80, 0.72, and 0.64, respectively), while the contribution of TGW was lower (R^2^ = 0.45). Nevertheless, as observed from significant *r* coefficients, the involvement of given traits to GYM2 formation in each environment was different. In line with linear regression analysis, the strongest and the most transversal positive correlation of GYM2 was observed with grain number and biomass, shown by a significant *r* value in eight out of the total nine environments tested (58-93% for GNM2 and 61-95% for BM2). In three low- and medium-yielding environments, GYM2 was positively influenced also by SNM2, GYS and GNSP (*r* = 50-79%). TGW confirmed to be less important for final grain yield, the two parameters being significantly correlated in only two medium-yielding environments (*r* = 60-82%). Finally, GNM2, the key trait for wheat grain yield increases, as expected, was primarily influenced by SNM2 (*r* = 62-96%, significant in five environments), grain number spike^−1^ (GNS) and spikelet^−1^ (GNSP) (*r* = 66-96%, significant in three and four environments, respectively), particularly in low- and medium-to-low environments.

**Fig. 2.**
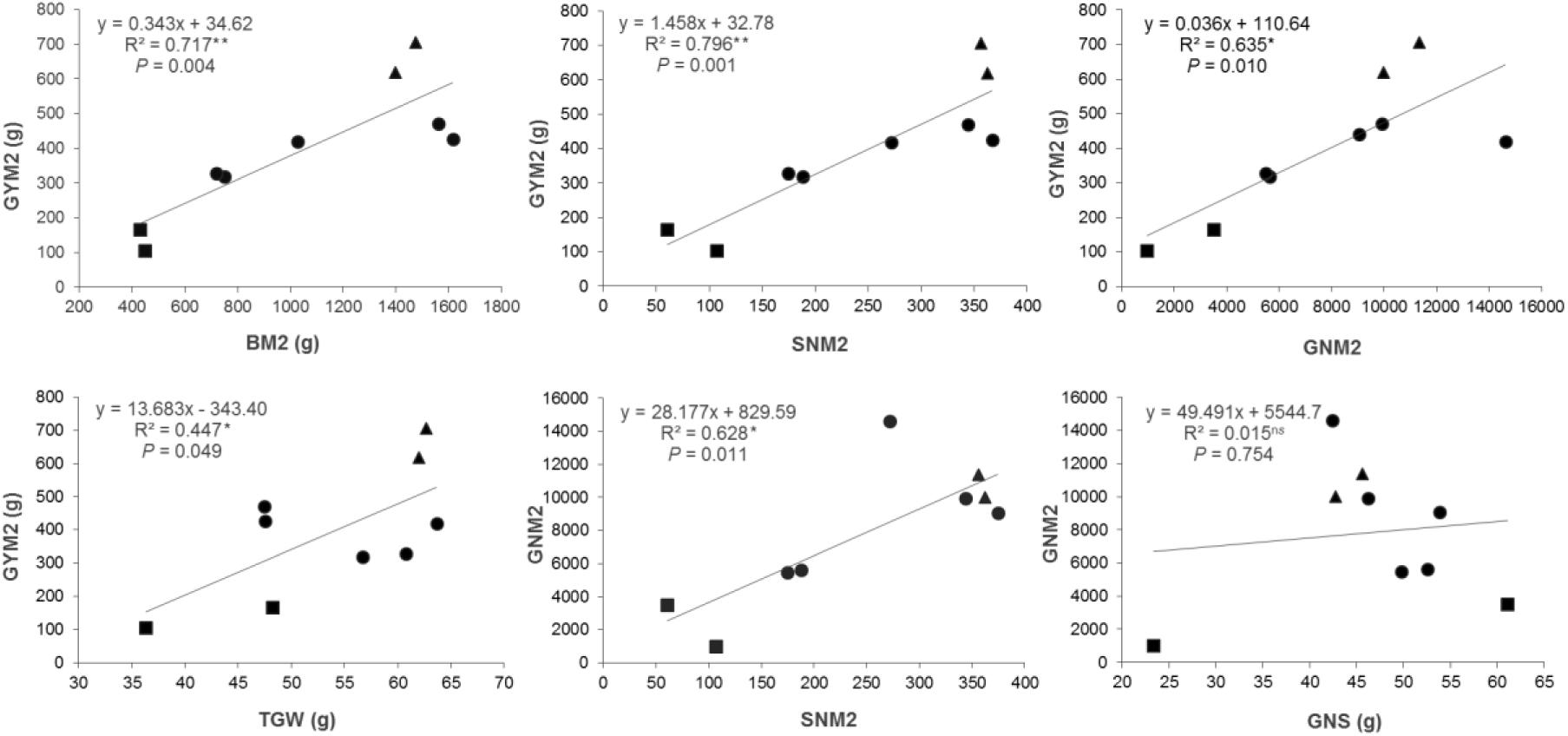
Scatterplots of the means of grain yield and grain number m^−2^ vs. main grain yield components from the three durum wheat-*Th. ponticum* NIRLs evaluated across nine environments (GYM2, grain yield m^−2^; BM2, biomass m^−2^; SNM2, spike number m^−2^; GNM2, grain number m^−2^; TGW, thousand-grain weight; GNS, grain number spike^−1^). \**P* <0.05, \*\**P*<0.01, “ns” not significant. Low- (GY < 2 t 374 ha^−1^), medium- (GY 2-5 t ha^−1^) and high- (GY > 5 t ha^−1^) yielding environments are represented by *square*, *circle* and *triangle* symbols, respectively.

**Table 6.**
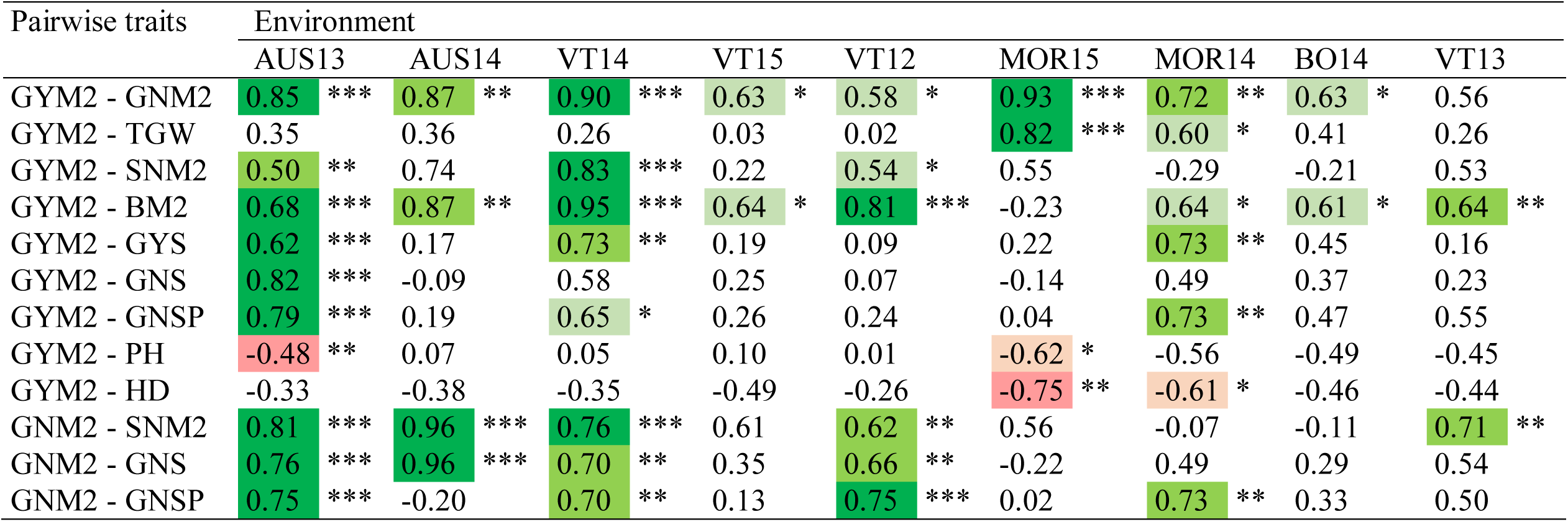
Pearson’s correlation coefficients between pairs of main yield-related traits involved in coarse and fine regulation of final grain yield across nine environments tested (GYM2, grain yield m^−2^; GNM2, grain number m^−2^; TGW, thousand grain weight; SNM2, spike number m^−2^; BM2, biomass m^−2^; GYS, grain yield spike^−1^; GNS, grain number spike^−1^; GNSP, grin number spikelet^−1^; PH, plant height; HD, heading date). Correlations are underlined using a heat colour map of green (positive correlations; dark green *P*<0.001, medium green *P*<0.01, light green *P*<0.05) and red (negative correlations; medium red *P*<0.01, light red *P*<0.05) shades. Environment acronyms are as described in Table 1.

| Pairwise traits | Environment |  |  |  |  |  |  |  |  |  |  |  |  |  |  |  |  |  |
| --- | --- | --- | --- | --- | --- | --- | --- | --- | --- | --- | --- | --- | --- | --- | --- | --- | --- | --- |
|  | AUS13 |  | AUS14 |  | VT14 |  | VT15 |  | VT12 |  | MOR15 |  | MOR14 |  | BO14 |  | VT13 |  |
| GYM2 - GNM2 | 0.85 | *** | 0.87 | ** | 0.90 | *** | 0.63 | * | 0.58 | * | 0.93 | *** | 0.72 | ** | 0.63 | * | 0.56 |  |
| GYM2 - TGW | 0.35 |  | 0.36 |  | 0.26 |  | 0.03 |  | 0.02 |  | 0.82 | *** | 0.60 | * | 0.41 |  | 0.26 |  |
| GYM2 - SNM2 | 0.50 | ** | 0.74 |  | 0.83 | *** | 0.22 |  | 0.54 | * | 0.55 |  | -0.29 |  | -0.21 |  | 0.53 |  |
| GYM2 - BM2 | 0.68 | *** | 0.87 | ** | 0.95 | *** | 0.64 | * | 0.81 | *** | -0.23 |  | 0.64 | * | 0.61 | * | 0.64 | ** |
| GYM2 - GYS | 0.62 | *** | 0.17 |  | 0.73 | ** | 0.19 |  | 0.09 |  | 0.22 |  | 0.73 | ** | 0.45 |  | 0.16 |  |
| GYM2 - GNS | 0.82 | *** | -0.09 |  | 0.58 |  | 0.25 |  | 0.07 |  | -0.14 |  | 0.49 |  | 0.37 |  | 0.23 |  |
| GYM2 - GNSP | 0.79 | *** | 0.19 |  | 0.65 | * | 0.26 |  | 0.24 |  | 0.04 |  | 0.73 | ** | 0.47 |  | 0.55 |  |
| GYM2 - PH | -0.48 | ** | 0.07 |  | 0.05 |  | 0.10 |  | 0.01 |  | -0.62 | * | -0.56 |  | -0.49 |  | -0.45 |  |
| GYM2 - HD | -0.33 |  | -0.38 |  | -0.35 |  | -0.49 |  | -0.26 |  | -0.75 | ** | -0.61 | * | -0.46 |  | -0.44 |  |
| GNM2 - SNM2 | 0.81 | *** | 0.96 | *** | 0.76 | *** | 0.61 |  | 0.62 | ** | 0.56 |  | -0.07 |  | -0.11 |  | 0.71 | ** |
| GNM2 - GNS | 0.76 | *** | 0.96 | *** | 0.70 | ** | 0.35 |  | 0.66 | ** | -0.22 |  | 0.49 |  | 0.29 |  | 0.54 |  |
| GNM2 - GNSP | 0.75 | *** | -0.20 |  | 0.70 | ** | 0.13 |  | 0.75 | *** | 0.02 |  | 0.73 | ** | 0.33 |  | 0.50 |  |

## Discussion

To evaluate, for the first time, the effect of different environmental conditions on yield and yield-contributing traits of three durum wheat-*Th. ponticum* recombinant lines, having 23%, 28% and 40% of 7AgL chromatin on their 7AL distal end, nine rain-fed field trials in four locations worldwide over four years were undertaken. The present analysis showed that the overall incidence of the 7AgL introgressions into the tetraploid wheat background on the trade-off between the various yield components, did not result, except for one environment (AUS14, see ahead), in major yield penalty, nor in any alteration of harvest index. In fact, taking all three 7AgL segments (all “+” vs. all “−” NIRLs comparison) and the nine environments together, an average, albeit not significant, 3% increase in grain yield m^−2^ was detected for the +NIRLs (Table 4). Given the high heterogeneity among environments, this minor difference resulted from four incremental cases (ranging from +1% to 18%) and five others exhibiting a yield reduction (1-2% in three of them). The same analysis of the effects of 7AgL segments as a whole but on individual yield-contributing traits highlighted that yield performances associated to the alien introductions were mainly due to significant increases of biomass m^−2^, spike/tiller number m^−2^, related grain number m^−2^ and spikelet^−1^, and to a general, significant decrease of grain weight (Table 4). This evidence confirms previous observations in one location only (Kuzmanović et al., 2014, 2016) and indicates stable expression of such traits across environments. The 7AgL case thus appears to support the consolidated evidence that in both bread and durum wheat, coarse regulation of yield is based primarily on grain number and biomass (Marti et al., 2016; Pedro et al., 2011; Slafer et al., 2014).

Grain yield of all NIRLs was generally higher in environments where more rainfall was recorded, particularly from heading onwards (e.g., Central and Northern Italy; Tables 1 and 2), which is in line with what observed for durum wheat grown under Mediterranean rain-fed conditions (Araus et al., 2003b). Similarly, previous studies indicate that bread wheat T4 derivatives, whose sizeable 7AgL segment includes those of the durum wheat recombinants described here, benefit from higher water availability (Singh et al., 1998; Monneveaux et al., 2003; Rosewarne et al., 2015). Nonetheless, the results from the environment-by-environment ANOVA (Table 4) revealed the potential of durum wheat 7AgL+ lines, taken as a whole, for yield increase also under heat and drought stress conditions, such as those of AUS13.

Considering the separate NIRL pairs across the range of environments, a number of 7AgL segment-associated effects emerged. R5+ exhibited an average 9% increase (albeit not significant) in grain yield (GYM2) vs. its control 7AgL− line (Table 5), as well as the highest grain yield gains with respect to the sites’ mean in eight out of the nine trials (Fig. 1b). While grain yield increase of R5+ vs. R5− plants was slight in high- and medium-yielding environments (1-4% in VT12, VT13, VT15, BO14, MOR15), in low-yielding AUS13 it amounted to a significant 52%, paralleled by a significant increase in several other yield parameters (Supplementary Table 3). These figures, however, were not mimicked by the AUS14 environment, which was characterized by similar meteorological conditions to AUS13 (Tables 1 and 2) but a longer growth cycle (earlier sowing) and a remarkable increase in flag leaf size of R5+ vs. R5− plants (Supplementary Table 3). Flag leaf size is known to be positively correlated with photosynthetic activity and yield in wheat under drought (Foulkes et al., 2007; Habash et al., 2007; Quarrie et al., 2006). The longer AUS14 growing season likely favoured vegetative plant growth, as also evidenced by greater biomass at anthesis (SDWA and TDWA) besides that bigger flag leaf area (Supplementary Table 3). However, the latter was probably not advantageous for R5+ (and so for R112+, see ahead), likely leading to high transpiration and dehydration rates, both negatively affecting final yield (e.g., Izanloo et al., 2008; Tardieu 2005).

As to the R112+ recombinant, average values of all trials showed increased values vs. R112− for spike (+14%) and grain number (+10%) m^−2^, as well as 6% higher grain yield (Table 5), in addition to a consistent tendency for wider flag leaf (significant in AUS14, VT12, VT13 and VT15, Supplementary Table 3) and higher chlorophyll content at late grain filling stages in several environments (Supplementary Table 3). These observations confirm that yield formation in R112+ depends mostly on tiller number development and potentially increased photosynthetic activity of leaves, both contributing to grain number formation (see also Kuzmanović et al., 2014, 2016). Tiller number is greatly influenced by the environment and determines wheat adaptive ability under rain-fed conditions (Elhani et al., 2007; Zhang et al., 2010); thus, it is not surprising that higher grain yield of R112+ vs. control plants (+3-7%) was observed in only three of the experimental sites, where more favourable conditions for tillering were evidently met (VT12, VT15, MOR15, Supplementary Table 3). At high-yielding sites, R112+ produced essentially the same as the controls (BO14, VT13), while a major yield penalty was recorded in low-yielding AUS14 (−29%, Supplementary Table 3), concomitantly with a significant, environment-specific decrease in spike fertility (−14% of grains spikelet^−1^). Similarly to the R5+ case (see above), the significantly larger flag leaf size (+25% FLW), accompanied by higher chlorophyll content (+23%, though not significant) at late grain filling in the R112+ case, seemed to be at a disadvantage for R112+ grain yield in AUS14 (Supplementary Table 3). Capacity of a wheat plant to maintain flag leaf greenness generally enhances photoassimilation (Foulkes et al., 2007; Peremarti et al., 2014); yet, in environments such as South Australia, where high irradiance often comes along with drought stress, increased temperatures and high wind (Izanloo et al., 2008; Fleury et al., 2010), high chlorophyll content could provoke oxidative damage and thus reduce photosynthetic capacity and, ultimately, yield potential (Long et al., 2015; Parry et al., 2011; Tricker et al., 2018; Zhu et al., 2010). Another possible reason for the reduced yield of R112+ in the Australian environment might reside in the interaction of its root system with the soil features at this site. Previous simulations determined that a shallow root system is more favourable in this type of environment (reviewed in Izanloo et al., 2008), which contrasts with the root system architecture (RSA) of R112+, found to be characterized by increased seminal root angle, total root length and root dry weight (Virili et al., 2015). Under drought conditions, a widened root angle and deeper roots were found to be crucial adaptive mechanisms associated to yield increases in rice (Uga et al., 2013; 2015; Ahmadi et al., 2014) and wheat (Lopes and Reynolds, 2010; Manchadi et al., 2006; Slack et al., 2018). However, the shallow, clay-limestone soils, typical of most Adelaide Plain’s and of our testing site, might have hindered the potential of the R112+ root system, limiting its access to water and nutrients at deeper layers. In fact, it was reported that root impedance reduces leaf elongation and the number of tillers in wheat (Jin et al., 2015). Therefore, the R112+ recombinant looks to be unsuitable for dry environments with combined stress factors, such as heat, drought and hostile soil structure. Instead, to exploit at best its 7AgL-linked positive effects on tiller/spike number, flag leaf photosynthetic activity (see also Kuzmanović et al., 2014, 2016), and RSA characteristics (Virili et al., 2015), it could be profitably employed in breeding directed to environments with optimal thermo-pluviometric patterns and soil characteristics (as VT12, VT15 and MOR15).

Finally, yield performance of the R23+ NIRL represents an intriguing case. In fact, while the background genotype, common to the R23− NIRL, was probably responsible for generally lower adaptability and consequent productivity throughout environments compared to the other two NIRL pairs (see § 3.2, Figure 1 and Supplementary Table 3), specific, both positive and negative, effects linked to its 40%-long 7AgL segment were consistently expressed in the multi-environment analysis (Table 5). The most important incremental effects validated throughout all trials were on spike fertility index, grain and spike number m^−2^ (Table 5 and Supplementary Table 3). Enhancement of the first two traits strongly supports the suggested existence of a large effect QTL for grain number within the 28-40% 7AgL portion specific to the R23+ introgression (Kuzmanović et al., 2014; 2016). Interestingly, a couple of important QTL for spike fertility traits (No. of spikelets and grains spikelet^−1^) were recently identified in the homoeologous 7AL region of durum wheat (Giunta et al., 2018). Furthermore, the results showed that R23+, similarly to R112+, and so in the shared 23-28% 7AgL segment, harbours a gene/QTL for tiller/spike number m^−2^, evidently a major factor of its yield potential, particularly in the dryer southern Australian (AUS13) and Moroccan (MOR14) environments (Supplementary Table 3; see also Kuzmanović et al., 2014; 2016). However, in contrast to the R112+ case, in which higher tiller number is likely to have primarily contributed to a more efficient plant source (i.e. leaf area, chlorophyll content), in R23+ the increased tiller number in combination with remarkably higher spike fertility had probably a more prominent effect on the genotype’s sink (i.e. grain number m^−2^). Confirming a different yield formation from that hypothesized for R112+, flag-leaf related traits were instead significantly depressed in R23+. Moreover, the same genotype also had reduced biomass at anthesis and maturity when compared to its R23− control (about −18%, Table 5). Both traits appear to be highly correlated with grain number (e.g. Fischer, 2008; González et al., 2011; Slafer et al., 2015); hence, it was unexpected that R23+ could consistently sustain its large gain in grain number m^−2^ and in spike fertility index (+15% and +46%, respectively) across eight environments (all but AUS14, where R23 was not tested; Table 5). Mechanisms involved in the control of fertile florets survival rate, rather than their absolute number, are likely to be responsible for this (Kuzmanović et al., 2016 and references therein). Recent work by Marti et al. (2016) established that growth and N partitioning in the two weeks before anthesis are key factors in determining differences in wheat yield performance, particularly for grain number. The same authors suggest that this growth period is not relevant for total biomass production at anthesis, but is crucial for determining the relative grain number. As the nature of this relationship is still uncertain (Slafer et al., 2015; Terrile et al., 2017 and references therein), it would be worthwhile investigating the modes of how assimilates are accumulated and translocated in R23+ during spike development just prior to anthesis. Guo et al. (2017) suggested a critical role of carbohydrate metabolism and phytohormones in regulating the floret primordia survival (FPS) and final grain number in wheat. Floret primordia formation (FPF) and their survival are known to be proportional to the availability of assimilates allocated for their development during spike growth before anthesis (Ferrante et al., 2013; González et al., 2011; Slafer et al., 2015). In R23+ plants, the number of fertile florets at anthesis (mirroring the FPF) showed to be similar to its R23- control (Kuzmanovic et al., 2016), while the number of seeds at maturity was enhanced, as a consequence of higher FPS (Table 5 and Supplementary Table 3; see also Kuzmanovic et al., 2014, 2016). Therefore, translocation of assimilates to grains is likely to be affected rather than their accumulation in spike tissues. Particularly noteworthy is the occurrence of increased spike fertility of R23+ vs. R23-not only in environments characterized by profuse rainfall throughout the crop cycle, such as VT14 and BO14, but also in AUS13 and MOR14, where these favourable conditions were not met (Tables 1 and 2; Supplementary Table 3). Despite the consistently lower thousand-grain weight of R23+ compared to R23- plants, higher grain number of R23+ was likely responsible for increments in grain yield (+7-58%) observed in four out of the eight environments (Table 5 and Supplementary Table 3). Due to the grain number vs. grain weight trade-off, increased fruiting efficiency is considered to be generally irrelevant for actual yield increases (Slafer et al., 2015). Nonetheless, this negative relationship may not be “constitutive”, i.e. the factor(s) increasing fruiting efficiency may be independent of the size of florets, i.e. potential grains (Slafer et al. 2015; Ferrante et al. 2015). Also, resource partitioning to developing florets may be increased and mortality of distal florets reduced, thus ensuring yield gains (Slafer et al., 2015, and references therein). Similar mechanisms may apply to the R23+ case. Interestingly, the highest grain yield increases of R23+ were observed in dry environments of AUS13 and MOR14 (+58% and +34%, respectively), indicating that R23+ maintains its higher seed set even under drought and/or heat stress conditions. This finding is of great relevance, as grain number is recognized as the most susceptible yield component to drought (Richards et al., 2010)., Furthermore, spike fertility was recently shown to be the key trait contributing to yield performance of durum wheat grown under severe and protracted heat stress (Sall et al., 2018).

The possibility of fully unlocking the R23+ yield potential for practical exploitation remains, however, somewhat challenging, because of the linkage between the gene(s)/QTL for grain number and spike fertility, and a gene(s) causing segregation distortion (*Sd*) and possibly correlated depression of a number of morpho-physiological traits, including grain development (Ceoloni et al., 2014b; Kuzmanović et al., 2016). Nonetheless, this linkage drag could be broken, e.g. via induced mutations or further homoeologous recombination, or possibly countered by transferring the R23 7AgL segment to different recipient cultivars, particularly choosing those exhibiting high grain weight, the trait mostly depressed in R23+ vs. R23- across environments (Table 5). In this view, previous evidence that, similar to the bread wheat T4 case, the interaction of the R23 40%-long 7AgL segment with varying durum wheat genetic backgrounds results in improved transmission of the recombinant chromosome, with little or no effect on plant phenotype (Ceoloni et al., 2014b), is encouraging.

Apart from genotype-dependent “stabilization” that the R23 7AgL segment may require, all segments can be readily moved into more site-directed (R112) or across environment (mainly R5 and R23) breeding pipelines for exploitation of their *Lr19+Sr25+Yp* genes, as well as for additional traits contributing to yield increase and stability. Moreover, all three recombinants can be further enriched with other useful alien genes. Recently, several new recombinants in bread and durum wheat have been obtained by chromosome engineering, in which highly effective gene(s)/QTL for resistance to *Fusarium* head blight and crown rot, originating from *Thinopyrum* species, were pyramided onto the most telomeric portions of the 7AgL segments described here (Ceoloni et al., 2017b; Forte et al., 2014; Kuzmanović et al., 2017). Preliminary results have shown normal fertility of the new recombinants and even higher yields when compared to control plants. This evidence contributes to strengthen the validity of targeted exploitation of alien variation to enhance yield potential in wheat species (Ceoloni et al., 2014a; Mondal et al., 2016; Zaïm et al., 2017, and references therein).

## Conclusions

Analysis of the three 7AgL introgression lines into durum wheat across a range of variable environments has validated their potential as donors of yield-contributing traits, besides that of additional beneficial attributes originating from the wild *Th. ponticum* donor. Being developed in a cultivar best adapted to Italian growing conditions (Simeto), the three recombinants exhibited variable performances in stressed environments, like South Australia and Morocco. In general, under no significant water stress, 7AgL lines responded well, without significant yield losses, mainly through contribution of grain and spike number, and, to some extent, flag leaf size and function at late grain filling stages. Also the observed significant increases in grain yield and grain number for 7AgL+ vs. 7AgL− NIRLs in more heat and drought-stressed environments, indicate that the 7AgL yield-related gene/QTL content may also be beneficial under adverse growing conditions. The analysed set of NIRLs represent a valuable toolkit for deciphering physiological mechanisms and identifying genes involved in grain yield regulation, as they display significantly different phenotypes for a number of traits associated with specific 7AgL segments.

## Acknowledgements

Financial support from MIUR (Italian Ministry of Education, University and Research), grant PRIN (Progetti di Ricerca scientifica di rilevante Interesse Nazionale) 2010–11 on ‘Identification and characterization of yield- and sustainability-related genes in durum wheat’, as well as from Lazio region—FILAS project “MIGLIORA”, are gratefully acknowledged. The authors would like to acknowledge Poh Chong and Alessandra Bitti for technical assistance with the Australian and Italian trials, respectively.

